# Extracting transition rates in single-particle tracking using analytical diffusion distribution analysis

**DOI:** 10.1101/2020.05.06.080176

**Authors:** Jochem N.A. Vink, Stan J.J. Brouns, Johannes Hohlbein

## Abstract

Single-particle tracking is an important technique in the life sciences to understand the kinetics of biomolecules. Observed diffusion coefficients *in vivo*, for example, enable researchers to determine whether biomolecules are moving alone, as part of a larger complex or are bound to large cellular components such as the membrane or chromosomal DNA. A remaining challenge has been to retrieve quantitative kinetic models especially for molecules that rapidly interchange between different diffusional states. Here, we present analytic diffusion distribution analysis (anaDDA), a framework that allows extracting transition rates from distributions of observed diffusion coefficients. We show that theoretically predicted distributions accurately match simulated distributions and that anaDDA outperforms existing methods to retrieve kinetics especially in the fast regime of 0.1-10 transitions per imaging frame. AnaDDA does account for the effects of confinement and tracking window boundaries. Furthermore, we added the option to perform global fitting of data acquired at different frame times, to allow complex models with multiple states to be fitted confidently. Previously, we have started to develop anaDDA to investigate the target search of CRISPR-Cas complexes. In this work, we have optimized the algorithms and reanalysed experimental data of DNA polymerase I diffusing in live *E. coli*. We found that long-lived DNA interaction by DNA polymerase are more abundant upon DNA damage, suggesting roles in DNA repair. We further revealed and quantified fast DNA probing interactions that last shorter than 10 ms. AnaDDA pushes the boundaries of the timescale of interactions that can be probed with single-particle tracking and is a mathematically rigorous framework that can be further expanded to extract detailed information about the behaviour of biomolecules in living cells.

## Introduction

Single-molecule studies have greatly expanded our knowledge of the mode of action and kinetics of DNA-protein interactions at the nanoscale^1^. Single-molecule Förster resonance energy transfer (smFRET) and optical/magnetic tweezers, for example, are well suited techniques to study forces, conformational changes and displacements of DNA-binding proteins such as DNA and RNA polymerases^2,3^, helicases^4,5^ and CRISPR-Cas proteins^6,7^ *in vitro* with high spatiotemporal resolution^8–11^. *In vivo*, however, single-particle tracking (SPT) remains the most convenient choice to study dynamic interactions^12^. For performing SPT, a gene of interest is fused to a gene expressing either a fluorescent protein or a protein tag (HaloTag/SnapTag) that can be later labelled with an organic fluorophore^13,14^. To avoid the temporal overlapping of emitters moving in the confined volume of (bacterial) cells, two strategies can be pursued. Either the expression level of the protein of interest is kept sufficiently low, or the emission signal from different proteins has to be separated in time which can be achieved using photoswitchable or photoactivatable fluorescent proteins or equivalent organic fluorophores enabling single-particle tracking photoactivation light microscopy (sptPALM)^15–18^. After linking subsequent localizations of these proteins into tracks, the apparent diffusion coefficient *D*\* is calculated from the mean squared displacement (MSD). The different mobilities of proteins switching between a DNA-bound state, in which proteins diffuse very slowly, and a DNA-free state, in which proteins diffuse through the cytoplasm, can provide kinetic information on the frequency and longevity of DNA-protein interactions.

The ability to extract this information, however, is compromised by photo bleaching, which limits the length of each track to a few localisations in the case of fluorescent proteins^19^. Furthermore, the limited localization precision increases the apparent diffusion of immobile states. Therefore, measured displacements cannot be unambiguously assigned to either a bound or a diffusing state. As a consequence, histograms of *D*\* values are often rather broad making a clear distinction between two diffusional states of a single species impossible. For the special case of non-interconverting *D*\* distributions, the shape of distributions can be calculated for a fixed number of analysed steps^20,21^ and, via fitting of the experimental data, used to extract the fractions of mobile and immobile proteins.

Another factor that can increase the overlap between two states in *D*\* distributions are state transitions occurring within single tracks. Using a typical frame time of 10 ms and a typical track length of 40 ms, any transition occurring within that track length will average out (Figure 1A). The framework described in references^20,21^ does not account for the possibility of transitions within a track. Consequently, the overlap can lead to overfitting, as an increase of intermediate values would necessitate the addition of more states, which are not necessarily biologically relevant. *In vitro* smFRET measurements have encountered a similar challenge, in which the interchanging of conformational states within single bursts or within single frames resulted in the averaging of FRET values. By implementing probability distribution analysis (PDA)^22,23^ previous studies were able to extract kinetic information and fit the entire FRET distribution^24–26^.

**Figure 1.**
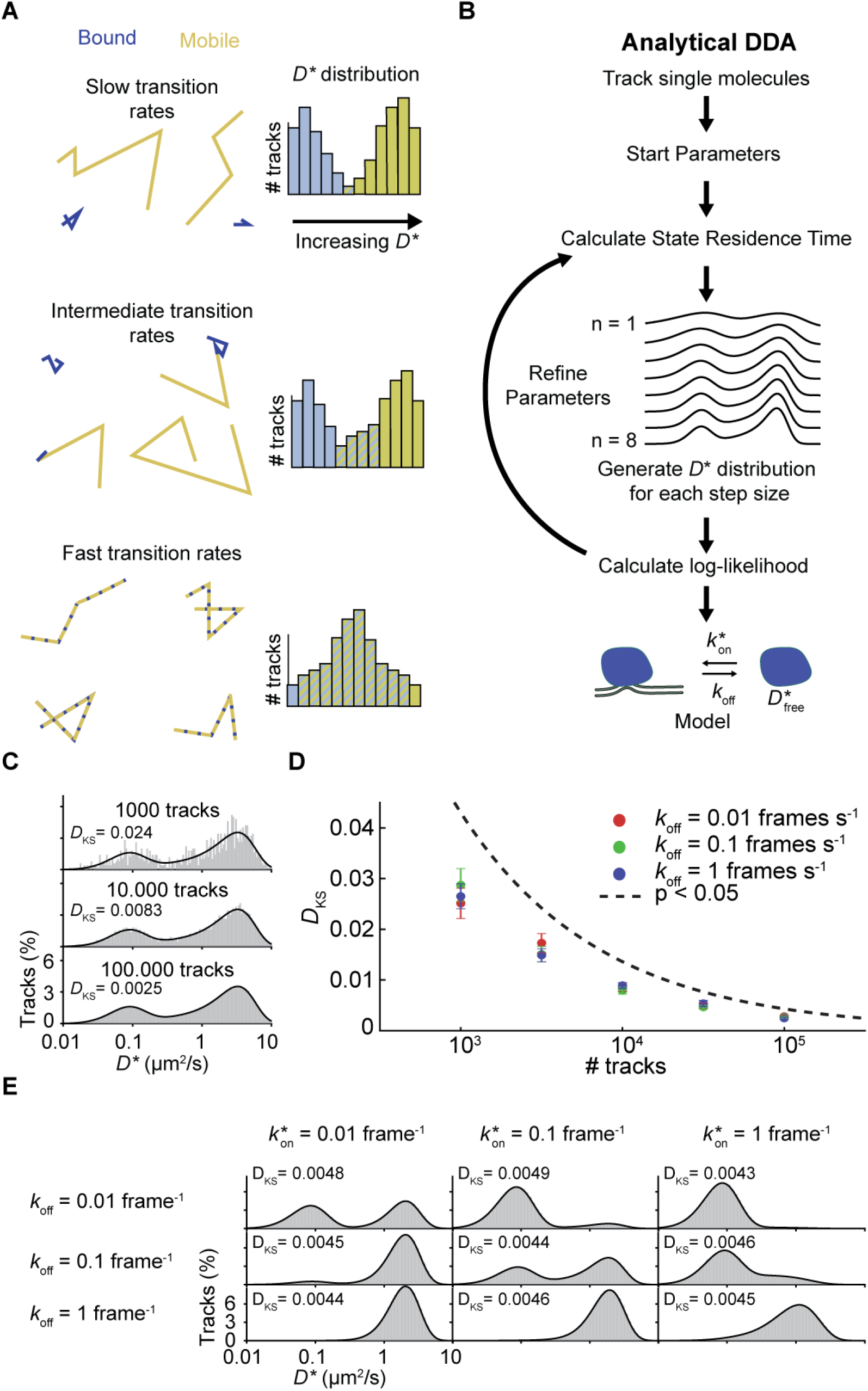
Analytical DDA. **(A)** The effect of transition rates on *D*\* distributions is depicted with simulated tracks of four steps and different transition rates. With increasing transition rates relative to the track length, the bound and unbound distributions start merging towards an intermediate apparent diffusion speed diffusivity (right). **(B)** Procedure of analytical DDA: The *D*\* values from tracked single particles are run into an MLE optimization program which refines a set of start parameters based on the likelihood to find a certain value given the number of tracks (all tracks longer than 8 are reduced to the first 8 steps). **(C)** Comparison of simulated (grey bars) and theoretically predicted (black line) distribution with different amount of tracks and the following starting parameters: 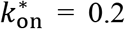 frame^−1^, 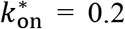 frame^−1^, *D*_free_ = 4 μm^2^/s and σ = 30 nm (localisation precision). Tracks are simulated without any confinement boundaries. The Kolmogorov-Smirnov test statistic (*D_KS_*) is indicated at each histogram. **(D)** The Kolmogorov-Smirnov test statistic compared to the threshold for statistically distinguishable distributions. Values above the threshold line indicate that two distributions significantly differ from each other. Error bars indicate S.E.M. of three independent simulations. **(E)** Comparison of simulated *D*\* distributions (grey) and the distributions calculated with analytical DDA for different transition rates (black). The shape of the distributions depends on both the ratio between 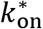 and *k*_off_ and the absolute values of these parameters. In this example *D*_free_ = 4 μm^2^/s and σ = 30 nm. For more tested parameters see Figure S1.

In this study, we aim to incorporate the statistical framework of PDA into *D*\* fitting of sptPALM data, which will allow us to directly extract biologically relevant parameters such as on- and off-rate next to the free diffusion coefficient and the total DNA-bound fraction. This method, which we call analytical diffusion distribution analysis (anaDDA), finds the kinetic parameters by implementing maximum likelihood estimation (MLE) and uses the probability to find *D*\* for all tracks, regardless of their length, present in the data set (Figure 1B). We benchmark this analysis method, with simulation of transitioning particles and implement modifications that account for specific experimental challenges, such as varying tracking windows and confinement effects within the cell. Furthermore, we compare anaDDA to a different kinetic analysis tools that use Bayesian statistics or unsupervised Gibbs sampling to infer state transitions from the data^27,28^. We study the effects of confinement and tracking parameters on the fitting of the distribution coefficient distribution and re-analyse previously published sptPALM data of DNA interacting proteins, obtain their kinetic parameters, and reveal that fast DNA probing interactions were hidden in the published data. Using anaDDA, we demonstrate the fast and accurate analysis of transient DNA-protein interactions using sptPALM in the millisecond time range, a range that was previously only accessible in slimfield microscopy^29^.

## Methods

### *D*\* fitting with transitioning states

Distributions of *D*\* have been fitted in numerous studies of DNA binding proteins^30,31^ using an formalism derived by Qian *et al*.^20^ from repeated convolution of the exponential distribution of displacement, resulting in a gamma function for each state. The formalism was later expanded by Michalet to account for localization errors^32^ leading to

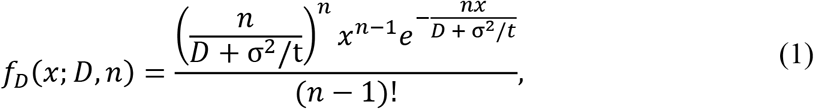

where *x* is the measured displacement, *D* is the apparent diffusion coefficient, *n* is the number of steps per track, *t* is the frame time and σ is the localization error. For multi-state (or multi-species) systems, terms can be added with different values of *D*_i_ and normalised by probability coefficients *A*_i_. The goal is to find the distribution of measured *D*\* values (*x*), for a certain number of underlying states that each have a probability *A*_i_ and a diffusion coefficient *D*_i_. These distributions assume, however, that there is no dynamic transitioning occurring between diffusional states of one species.

In order to account for dynamics of state transitions in a two state system, we incorporated a statistical framework derived for probability distribution analysis (PDA) that is used to analyse single-molecule FRET distributions^22,23,33^. This method describes the distribution of time spent in each state given a certain 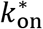, *k*_off_ and the integrated time *t*_int_.

Firstly, the probability distribution function after starting from state S1 and for a time interval *t*_int_ can be calculated by three equations corresponding to 0, an odd or an even number of transitions^23^:

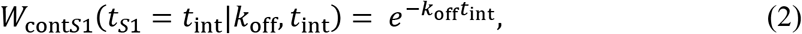

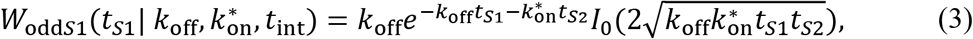

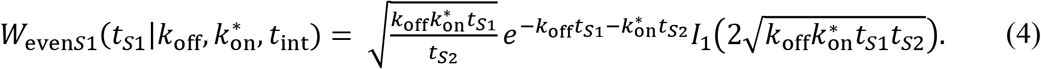

Where *t*_*S*1_ and *t*_*S*2_ are times spent in state *S*1 and state *S*2 and *I*_0_ and *I*_1_ are Bessel functions of order zero and one, respectively. Note that *t*_*S*1_ + *t*_*S*2_ = *t*_int_. Equations for starting in state *S*2 (*W*_contS2_, *W*_oddS2_ and *W*_evenS2_), can be found by exchanging *k*_off_ for 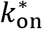 and *t*_*S*1_ for *t*_*S*2_ and vice versa in equations 2–4.

To correctly describe the distribution over a certain number of frames, we first calculated the distribution over a single time frame *t_f_*. Within a single frame, a particle started in that state can either end in the same state or in a different state. Therefore, in a two-state system the probability function for four scenarios have to be calculated

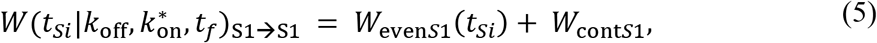

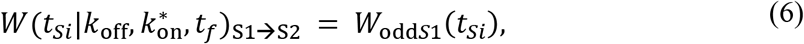

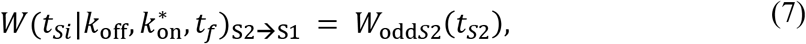

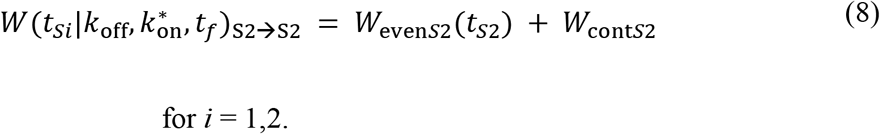

To link the distribution of time spent in a state to the distribution of measured displacements (*x*), we can convert the time spent in each state and its diffusion coefficient to the average observed diffusion coefficient by the following equation (assuming here that *S*1 is an immobile state)

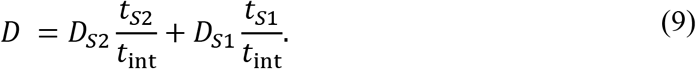

In case of the transition between an immobile bound state *S*1 (*D*_*S*1_= 0) and a mobile state with diffusion coefficient *D*_*S*2_ = *D*_free_ we can modify the above equation to

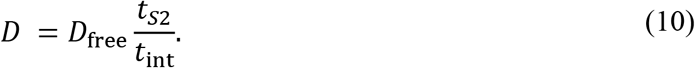

For all further work we assume that the first state is immobile and use equation 10.

Using equation 10, the probability distribution function (equation 1) can be modified according to

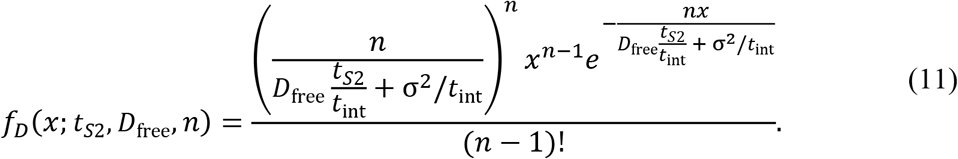

Subsequently, the probability to find a certain diffusion coefficient (*x*) for a single time step given the time spent in the mobile state is given by *f_D_*(*x*|*t*_*S*2_, 1). We can then find the distribution of measured diffusion coefficients for a single frame by integrating over all possible times spent in the mobile state

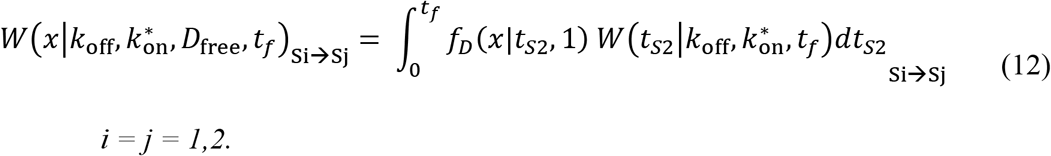

Now that we have the distribution for a single time step, we need to find the distribution for the average of multiple frames. For this we use the same method as Qian *et al*.^20^, namely repeated convolution of the distribution for a single frame, while keeping track of the start and end state. The probability distributions are therefore

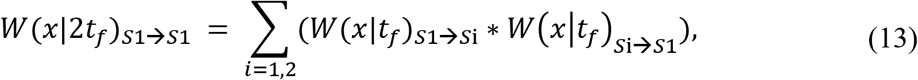

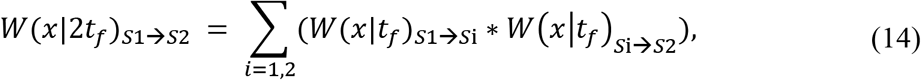

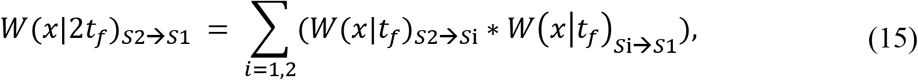

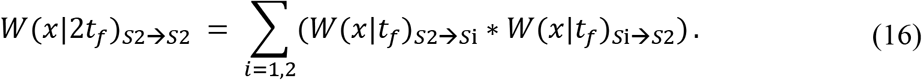

For a track consisting of 4 frames, the distributions found for 2 frames can be convoluted again. The full distribution is then found by summing up each of the partial distributions multiplied by the chance they start in *S*1 or *S*2:

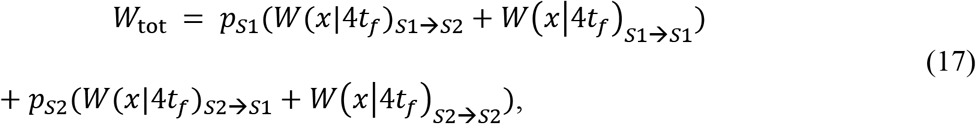

with *p*_*S*1_ and *p*_*S*2_ defined in equations 18 and 19, respectively:

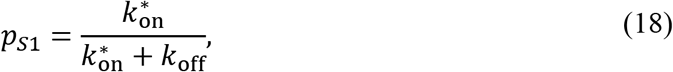

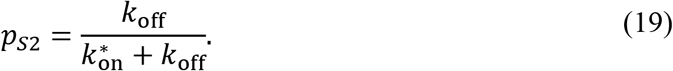

### Localization error

As two consecutive steps share at least one localisation, the localisation error of this localisation leads to a correlation between the step lengths^32^. Only in the special case of the localisation error being zero, the step lengths are uncorrelated. The distribution of the sum of step lengths for a certain number of steps is therefore not described by a gamma distribution, which is the sum of independent variables. However, as each step separately is a gamma random variable, we require the summation of correlated gamma random variables to describe the distribution of localization error analytically for different amount of time steps.

The extend by which the localization error affects the correlation of sequential steps can be quantified by calculating the correlation coefficient *ρ_i_* = 〈*x*,*y*〉/*σ_x_σ_y_* and the covariance of sequential steps as derived by Berglund^34^

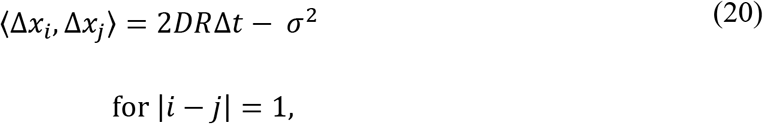

where *R* is the motion blur coefficient caused by movement of the particle during the illumination time and *D* is the diffusion coefficient. We assume further that measurements were taken with very short illumination pulses leading to *R*→ 0. We further convert equation 20 to MSD notation

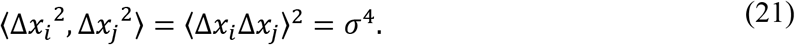

After converting to two dimensions and assuming that Δ*x* and Δ*y* are independent, we get

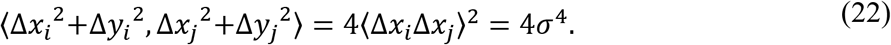

To calculate the correlation coefficient *ρ_ij_*, we use the following expression for the standard deviation of the MSD in two dimensions^32^

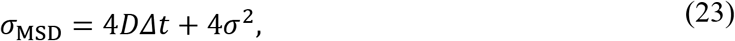

leading to

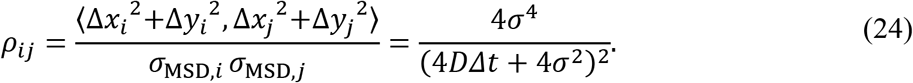

For most applications *DΔt* > *σ*^2^ and *ρ* can be neglected. However, for immobile particles *DΔt* = 0, and 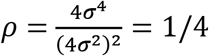. For a number of *n* measured steps with localization error of an immobile particle, the correlation matrix *ρ* is therefore given by

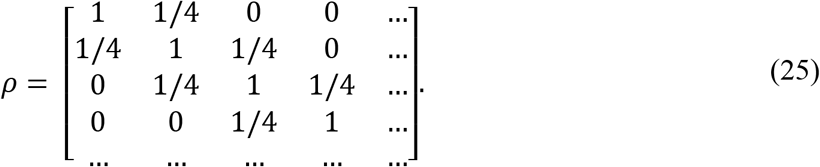

The summation of gamma random variables given a certain correlation matrix has been previously derived in terms of confluent Lauricella series^35^. Using the definitions above, this equation can be written as

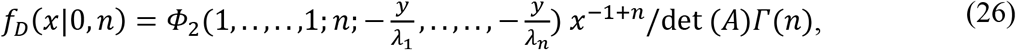

where *Φ*_2_ is the confluent Lauricella function, *λ*_1_-*λ_n_* are the eigenvalues of the matrix *A* = *B* · *B*, where *B* is an *n* x *n* matrix with diagonal values *σ*^2^, and *C* is an *n* x *n* matrix with values 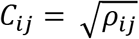.

This summation, for each number of measured steps *n* is the modified distribution for immobile particles taking into account the correlation between sequential measured displacements. To implement this distribution in the calculation of our total *D* distributions, we subtract the fraction of immobile particles after *n* time steps (*W*_*contS*1_(*t*_*S*1_ = 4*t_f_*), Eq.5) multiplied with the distribution of expected *D*\* for *n* time steps *f_D_*(*x*|0, *n*) (Eq. 1) and replace it with the same fraction of immobilized particles multiplied with the distribution calculated based on the Lauricella series. The calculation of confluent Lauricella series was implemented from MATLAB code described in Martos-Naya et al. (2016)^36^.

The equation above can be further refined to experimental data, if there is a large difference in localization error between particles. In that case, there is another correlation factor due to the difference in brightness/focus of particles. As some particles might show a dynamic brightness, e.g. by diffusing in and out the excitation/detection focus, localizations of this track will have a higher precision the brighter the emission of the particle is, altering the correlation matrix to

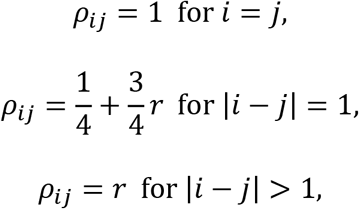

where *r* is the correlation coefficient between two steps within the same track not sharing any localizations (|*i* – *j*| > 1). We found that this correlation coefficient can be experimentally determined by measuring correlation of displacement of immobilized particles, or by measuring the correlation of estimated localization errors within tracks. This can be done by making a matrix in MATLAB where the rows are the different tracks and the columns are either the different step size of immobilized particles or the estimated localization errors. The built-in function *coerrcoef* then automatically calculates the correlation coefficient of this dataset.

### Tracking window

In order to the prevent the accidental linking of different diffusing particles, many tracking algorithms use a certain cut-off, in which steps longer than a certain distance are not allowed^37–39^. However, this tracking window can influence the distribution of *D* values recovered. In analytical DDA, we correct for this by setting *f_D_*(*x* > *maxD*|*D_i_*, 1) = 0, where *maxD* is the maximum *D*\* value that can be obtained given the tracking window.

### Confinement

To take the effects of geometrical confinement within the cell into account, we implemented an analytical way to calculate the effective diffusion coefficient given the geometry and the real diffusion coefficient. Most boundary geometries encountered in *in vivo* settings are either spherical or rod-shaped. For a spherical geometry, the effective measured MSD given a diffusion coefficient *D* and a timestep Δ*t* have been previously derived^40^. First, the authors defined the zeros *α_m_* at which 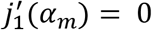, with 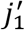 being the derivative of the spherical Bessel function of the first kind. This can be rewritten as

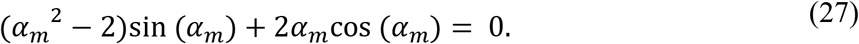

Subsequently the effective measured MSD within a spherical confined space of radius *r* is equal to

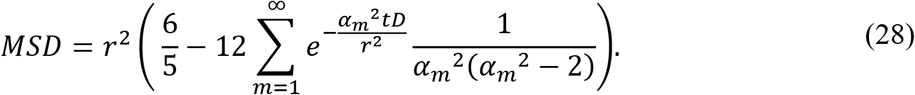

This infinite series converges to zero. We therefore used the first 10.000 terms for calculation as a reasonable approximation. Because the previous equation refers to the three-dimensional MSD we use the following relation to calculate the observed diffusion coefficient we divide by 6*t*,

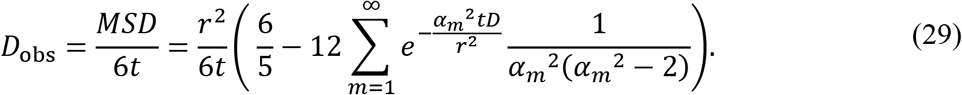

We can define the above equation as a function to calculate the observed diffusion given a certain radius, frame time and diffusion coefficient in the presence of spherical confinement

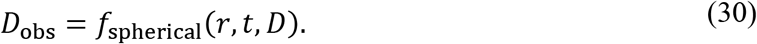

We then substitute *f_D_*(*x*|*D*, 1) for *f_D_*(*x*|*D*_obs_, 1) and use equation 12 to calculate the distribution under any number of steps and given a *k*_off_ and 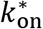.

For rod geometries, there is no analytically derived solution available. However, we can combine the spherical derivation with a derivation in the same study for circular 2D geometries. In this geometry, the authors defined zeros *β_m_* of the function 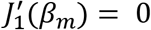, where 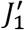 is the derivative of the Bessel function of the first kind. Subsequently, the effective measured MSD within a circular confined space of radius *r* is equal to

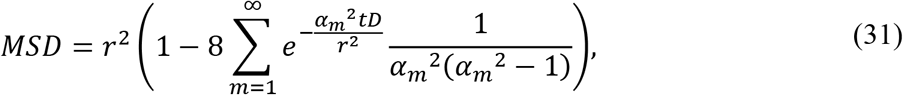

which we can again convert to a function to calculate the observed diffusion coefficient, but now as the MSD is two-dimensional, we divide by 4*t*

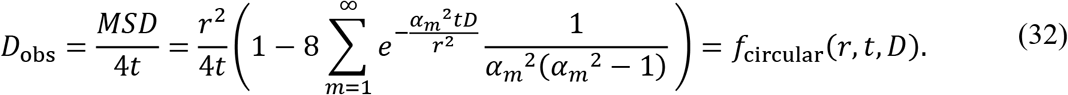

To calculate the effective measured MSD in a rod-shaped geometry, we split the cell in two parts: the hemispherical (consisting of two hemi-spheres) and the cylindrical part. If the cell is much longer than it is wide the cylindrical part dominates. For diffusion within a cylinder, movement along the cell length is not restricted, whereas movement along the width of the cell is constrained as given by equation 31. If the cell is as long as wide, we have a spherical cell for which the diffusion is described by equation 28. For cells featuring intermediate aspect ratios, we can calculate the ratio of these two domains via the ratio of their volumes

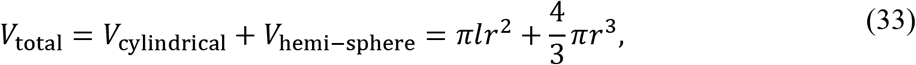

where *r* is the radius of the cell width and *l* the length of the cylindrical part of the rod-shaped cell. The observed diffusion coefficient along the cell length, *D*_obs,x_ is not being restricted in the cylindrical part. The observed diffusion coefficient along the cell width *D*_obs,y_ on the contrary is restricted within the cylindrical part. Therefore, we separately calculate these two observed diffusion coefficients

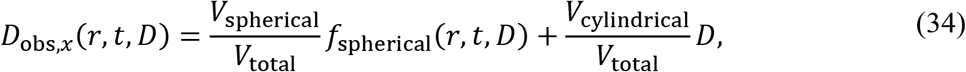

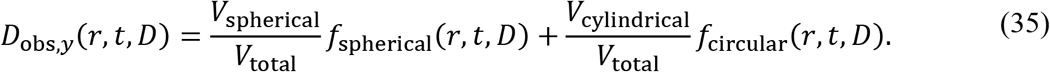

For the last step, we require a probability distribution function of the sum of the two distributions *D*_obs,*x*_ and *D*_obs,*y*_ and go back to the distribution of *x*

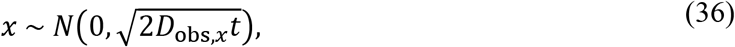

representing a normal distribution with mean of zero and 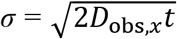. The distribution of the squared displacement is therefore a chi-square distribution

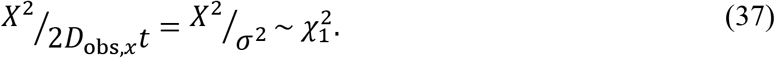

The same holds for *y* with

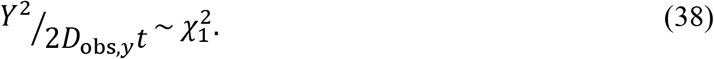

To get to the distribution of *D*_obs_ we calculate

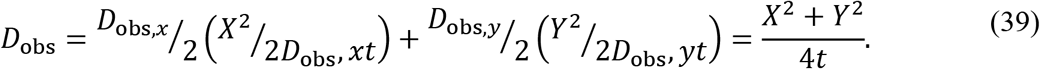

Consequently, the distribution of *D*_obs_ is a summation of two chi-square variables weighted by the different diffusion coefficients. The formula for this summation was given in a previous study^41^ for the following case: Let *X*, 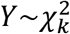 two independent and identically distributed chi-square random variables with *k* degrees of freedom. Let *Z* := *aX* + *bY*, then the density function *f_z_* is given by:

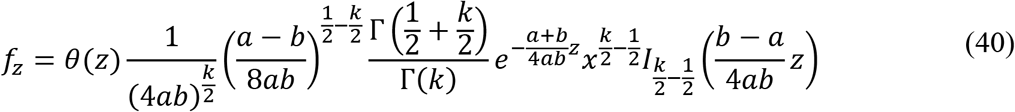

In our case, where *k* = *1*, this equation is simplified to

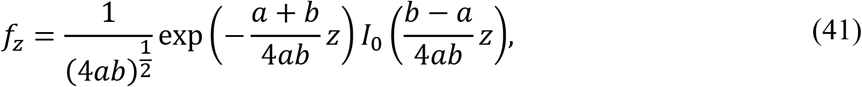

where *I*_0_ is the zeroth order modified Bessel function of the first kind. When we substitute ^*D*_obs,*x*_^/_2_ and ^*D*_obs,*y*_^/_2_ for *a* and *b* respectively (combine equation 39 and 41), we obtain the following equation for the summation of two diffusion coefficients in two dimensions

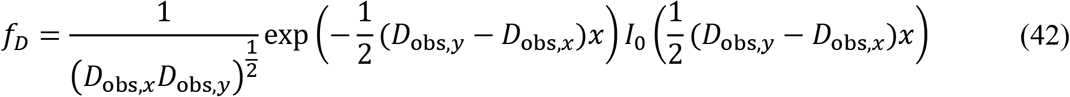

where *x* is the measured displacement as in equation 1. This distribution can then be used as substitution for *f_D_*(*x*|*D*, 1) and we can use equation 12 to solve the distribution under any number of steps and given a *k*_off_ and 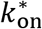.

### Maximum likelihood estimation (MLE)

To find the underlying parameters of experimental data and simulations, we use MLE which maximizes the joint probability of observing by iteration through the parameter space. Generally, MLE requires a probability density function to calculate and sum all probabilities of each observed data point. The benefit of the method is that it does not require any binning, compared to other optimization methods. However, MLE does require the exact probability for each data point to be calculable. Because we use numerical convolution (for increasing the performance of the algorithm, we implemented an FFT convolution^42^), we will only get the probability at discrete points within the probability density function. Therefore, to calculate the probabilities for the points of our data set, we use spline interpolation.

Because MLE is known to be affected by local minima^43^, we use a number of cycles (generally four) in which we generate random starting parameters and run the algorithm several times after which we select the end parameter set with the maximal likelihood. Those parameters are then used as starting parameters for bootstrapping in which we run the analysis through a number of subsets of the data to get an estimate of the standard deviations of our parameter estimates.

### Plotting of diffusion distribution histograms

With the parameter sets used in our simulations, the diffusion histograms are visually more distinguishable when log(*D*\*) is plotted compared to *D*\*. We therefore integrated the linear density function with widths specified by the bin size of the logarithmic scale to calculate the probability density function for log(*D*\*) instead of *D*\*.

## Results

### AnaDDA generates *D*\* distributions equal to the ground truth of simulated distributions

AnaDDA allows calculating the shape of the *D*\* distribution, depending on the free diffusion coefficient (the diffusion coefficient in absence of binding interactions) and the transition rates. As this shape depends on the step length of the tracks, we separate the tracks according to their respective length and fit each data point to the distribution that matches their step length. To benchmark our new analysis method, we first compared our theoretical predictions of the *D*\* distribution to data in which we simulated the diffusional characteristics of a particle that dynamically switches between a (DNA-) bound state and a freely diffusing state without including any boundary conditions for diffusion (see section below for confinement within cells). With increasing number of tracks, the predicted *D*\* distribution increasingly resembles the predicted theoretical distribution (Figure 1C). To test whether our theoretical distributions differed from the simulated ground truth, we performed Kolmogorov-Smirnov tests. We found that the test statistic **D_KS_** converged to zero for larger number of tracks analysed and was on average smaller than the critical value required to reject the null-hypothesis (*D_KS_* = 0.004 for p < 0.05), indicating that the ground-truth simulations and our theoretical predictions come from the same distribution (Figure 1D).

We varied the range of transition timescales (Figure 1E) ranging from 0.01 to 10 transitions per frame (at 0.01 s frame time) at all different step lengths included in this analysis (1-8; Figure S1) and compared a range of frame times (20-100 Hz) and experimentally realistic localization errors (20-50 nm) (Figure S2). Under all these conditions, the ground truth simulations (N = 100.000 tracks) and the anaDDA generated distributions showed very close agreement (*D_KS_* < 0.004). As this analysis involved a direct comparison between the predicted and simulated distribution without fitting the data or any optimization of parameters, it can be concluded that our theoretically predicted distributions are similar to the ground truth distributions.

### AnaDDA can extract transition rates from tracks with more than one transition per frame

With data from experimental measurements, the ground truth is unknown, and parameters have to be inferred by fitting. First, we tested via simulations how reliably parameters can be extracted over a large dynamic range of transitions. We compared the input parameters to the extracted ones with Maximum Likelihood Estimation (MLE). To benchmark the performance of extraction we calculate the accuracy through the geometric mean and the precision through the geometric standard deviation of 10 independent simulations. For all tested data sizes (5000-100000 tracks) and transition rates (0.001-10 transitions per frame), the analysis method is accurate (< ± 5% of input parameters). The precision decreased slightly with decreasing data size and for small/large transition rates (Figure 2). Furthermore, the precision at high transition rates (>1 transition per frame) is lower for 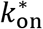 than *k*_off_ (Figure 2A-B). In general, the highest precision is found for tracks between 0.1 and 1 transition per frame. With 50.000 tracks per simulation, the transition rates over three orders of magnitude (0.002-2 transitions per frame) were determined with an error smaller than 20% of the actual value (Figure 2A-C).

**Figure 2.**
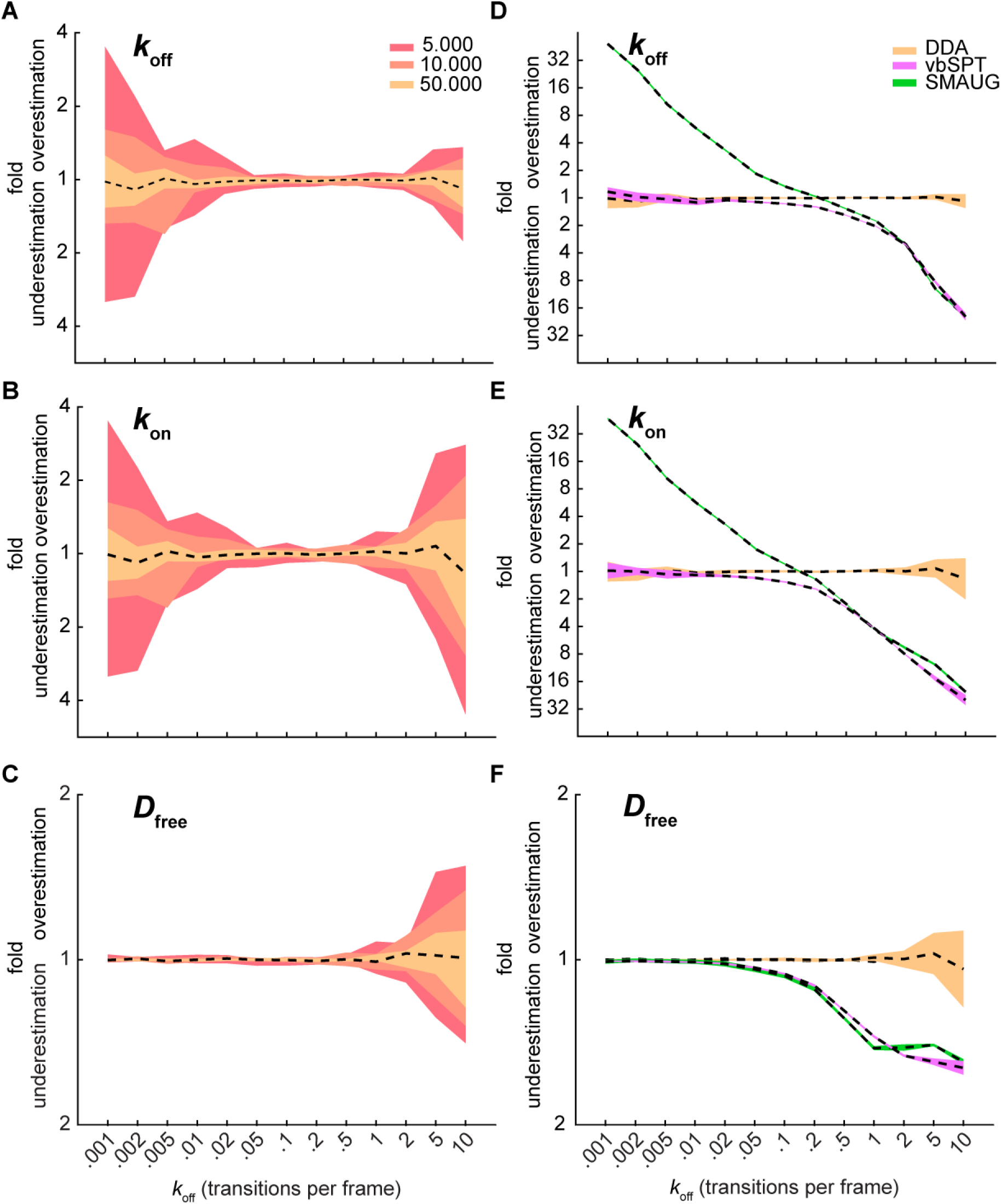
MLE extraction of parameters. The accuracy is calculated through the value of the geometric mean (dashed black line) and the precision is calculated through the geometric standard deviation of 10 independent simulations. The length of tracks was exponentially distributed with a mean of three steps and a cut off at 8 steps (*D*_free_ = 4 μm^2^/s, σ = 30 nm). **(A-C)** Effect of data size on accuracy and precision of extraction of (A) *k*_off_, (B) 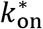 and (C) *D*_free_ for n=5.000 tracks (yellow), 10.000 tracks (orange) and 50.000 tracks (red). **(D-F)** Comparison of anaDDA versus vbSPT and SMAUG on accuracy and precision of extraction of (D) *k*_off_, (E) 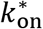 and (F) *D*_free_. 50.000 tracks were used for both methods.

We compared our method with a previously published framework that used Bayesian statistics to infer transition and diffusion dynamics (vbSPT)^27^ and a framework that used unsupervised Gibbs sampling for similar purposes (SMAUG)^28^. As vbSPT and SMAUG deduce the number of states from the data, we limited the amount of states in this analysis software to two to achieve a fair comparison. For slow transitions (<0.01 transition per frame) both anaDDA and vbSPT were able to extract the correct kinetic parameters (<20% error; Figure 2D-F), whereas SMAUG overestimated the transition rates. At faster transitions (> 0.02 transitions per frame), however, we observed a decrease in the extracted apparent free diffusion coefficient and a decrease in the extracted on and off-rates for both vbSPT and SMAUG.

When we removed the restriction of a two-state model, vbSPT started introducing multiple false states (Figure S3). Already at low transition rates (0.01 transitions per frame), vbSPT suggests the presence of a false third state. At this transition rate, two states (0.06 and 0.11 μm^2^/s) were close to the expected average apparent diffusion coefficient of the simulated immobile state (σ^2^/t = 0.09 μm^2^/s). The highest number of predicted states (4 states) was found for transition rates between 0.05 and 0.5 transitions per frame.

Our findings suggest that vbSPT and SMAUG fail to account for the increasing occurrence of multiple transitions within a single frame at fast transition rates. Our analysis software is distinctive in its ability to extract kinetic parameters when multiple transitions are likely to occur within the time window of the measurement. We wanted to further correct for artefacts that can influence diffusion distribution analysis, namely confined diffusion within cells and application of tracking windows.

### AnaDDA corrects for confinement within cells and restricted tracking windows

To study the effect of geometrical confinement, we simulated diffusive particles within the confined boundaries of different cell shapes. We previously showed that confinement only has a very small effect on observed transition rates in bacterial cells^44^. However, as the measured diffusion coefficient can be greatly affected by confinement, we implemented an algorithm based on previously developed derivations^40^ (for details see Materials and Methods) to account for confinement in both rod-shaped (e.g. *E. coli* cells) and spherical-shaped boundaries (e.g. eukaryotic nuclei) (Figure 3A).

**Figure 3.**
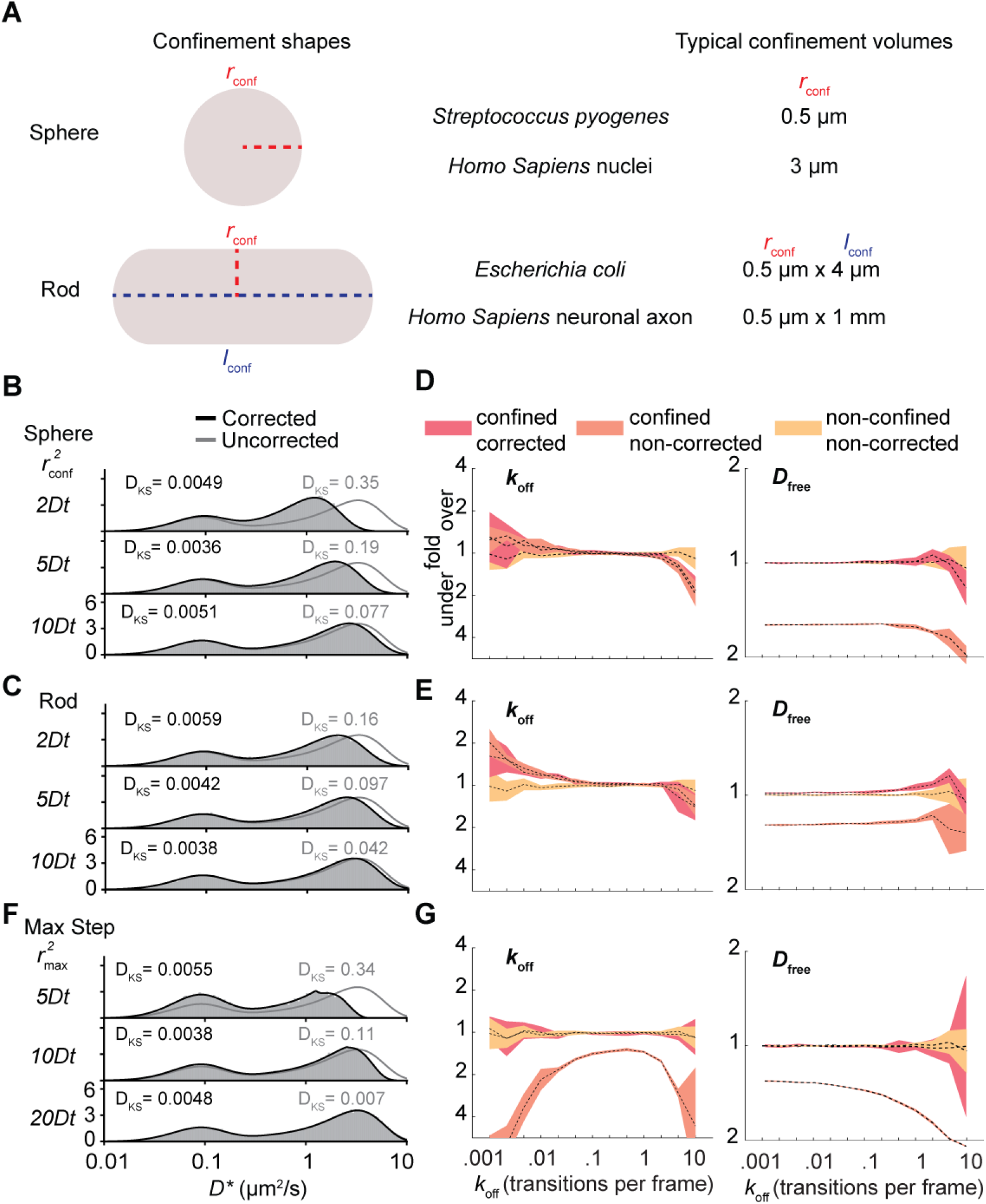
Effects of geometrical confinement and the length of the tracking window. **(A)** Typical confinement shapes within cells. The boundary shape of spherical cells is defined by a single parameter (radius; *r*_conf._), whereas rod-shaped cells are defined by two parameters (radius and length; *r*_conf._ and *l*_conf._). **(B-C)** Influence of spherical and rod-shaped boundaries on the distribution of simulated (grey box) and uncorrected DDA (grey line) and corrected DDA (black line) distributions. (*k*_off_ = 0.2 frame^−1^, 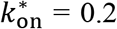 frame^−1^) **(D-E)** Influence of spherical and rod-shaped cells on the estimation of parameters of DDA on unconfined simulated trajectories (yellow), uncorrected DDA on confined simulated trajectories (orange) and corrected DDA on confined trajectories (red). **(F-G)** Same as B-E except for simulated trajectories with a maximum step size. Simulation parameters: *D*_free_ = 4 μm^2^/s and σ = 30 nm (localisation precision), N = 50.000. The Kolmogorov-Smirnov test statistic (*D_KS_*) is indicated in each histogram.

For both spherical and rod-shaped cells (cell length : radius = 8:1) we found that our theoretical predictions for varying cell sizes (*r*^2^ = (2,5, or 20)*D*_free_*t*) match well with simulated data (Figure 3B-C; *D_KS_* < 0.006) in contrast to uncorrected distributions for which the predicted distributions are statistically different from the simulated distributions (*D_KS_* > 0.04). In an *E. coli* cell (*r* = 0.5 μm) and under standard measurement frame times (0.01 s), these confinement regimes (*D*_free_*t*) would be reached with *D*_free_ values of 12.5 μm^2^/s respectively, which matches the values found for small single fluorescent proteins ^45^. In a eukaryotic nucleus (*r* = 3 μm), these regimes would correspond to *D*_free_ values up to 450 μm^2^/s which is generally much faster than any reported literature values. This finding indicates that geometrical confinement by cell boundaries is mostly limiting in prokaryotic studies. However, at longer frame times (0.1 s) confinement effects will play a role when studying diffusion within eukaryotic nuclei.

As not every cell in a population is the same size, the distribution might be further affected by a variation of cell sizes. We therefore analysed a mixture of three different simulated cell sizes and found that the distributions remained statistically indistinguishable from a uniform population of the same cell size (Fig. S4; *D_KS_* < 0.006). This shows that the correction method remains valid as long as the average dimensions of the cell boundaries are known.

To further test our ability to infer parameters from the data in a system where diffusion is geometrically confined, we performed MLE with and without corrections for confinement. We observe that the incorporation of our confinement corrections increases the accuracy and precision of the estimation of *D*_free_ (Fig. 3D-E). Compared to unconfined diffusion, there is a bias in recovered transition rates at very small and large transition rates, as these regimes are most sensitive to small deviations of the predicted distribution to the ground truth. These minor deviations are most likely caused by a correlation which occurs for diffusing particles within boundaries, where particles that are close to the boundary in one frame, are again likely to encounter the boundary in the next frame. That effect is not taken into account in our current implementation. However, for most transition regimes (0.01-2 transitions per frame), the error of the estimated parameters falls within 20%.

Another type of analysis artefact comes from the settings for tracking windows. When the density of labelled fluorophores is higher than 1 per cell, different molecules can be falsely assigned to the same track. To prevent this effect, multiple tracking software algorithms set a limit to the maximum step length that individual tracks are allowed to have. Although this is sometimes unavoidable, the absence of the largest steps can severely affect the MLE fitting parameters. AnaDDA is able to correct for this, by integrating this max step in the probability distribution (see Materials and Methods). The effect of this correction was tested for a range of radii of tracking windows (*r*^2^ = (5,10, or 20)*D*_free_*t*) and in all cases the *D_KS_* of the corrected distributions were below the threshold for significantly different distributions (*D_KS_* = 0.006), whereas for small and intermediate tracking windows (*r*^2^ = (5 and 10)*D*_free_*t*) uncorrected distributions were significantly different (*D_KS_* = 0.34 and *D_KS_* = 0.11; Figure 3F-G). The tracking window also had a large effect on both the predicted transition rates and free diffusion coefficients from MLE, where in the absence of corrections all parameters were significantly underestimated (>1.5x). With the correction, the estimations were again unbiased and very similar to the accuracy and precision of estimations in the absence of tracking windows.

Taken together, anaDDA can correct the distributions for measurements that are affected by confinement within spherical and rod-shaped boundaries and by the application of a maximum step size within tracking algorithms.

### AnaDDA can be expanded for multiple states and can integrate multiple frame times

So far, we have discussed the presence of one diffusing species converting between two diffusional states. In the following, we will expand the DDA-fitting to account for more species and states. Many DNA binding proteins contain both non- and target-specific interactions with DNA. Therefore, it is likely that the kinetics of these two interactions are different, which would require the model to be expanded beyond a two-state model. PDA statistical analysis currently does not incorporate more than two dynamic states. However, it is possible to incorporate more states by assuming that their dynamics are much slower than the non-specific DNA interactions, which would result in a negligible amount of transitions in the timeframe studied. Then these states can be approximated by separate static (non-interchanging) species (Figure 4A). Generally the specific interactions are much longer lasting than the non-specific interactions^46^, so in many cases this assumption would be valid.

**Figure 4.**
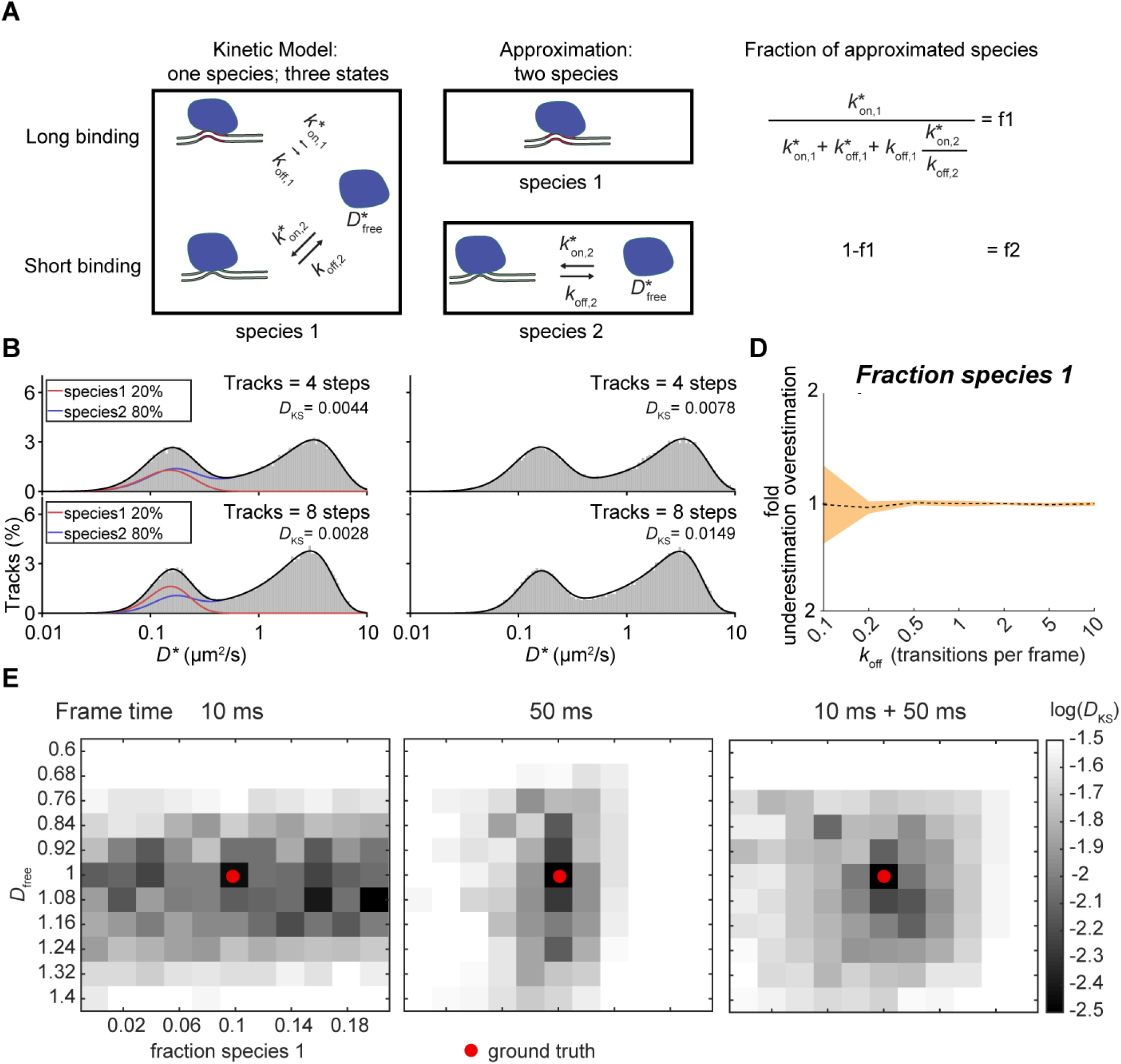
Three-state models and multiple frame times. **(A)** Three-state models cannot be directly described with PDA statistics. If some interactions are slower than the typical frame time, however, the approximation can be made that they belong to a non-transitioning separate species. The expected fraction of each of this species can be calculated from the on and off-rates of all states (right). **(B)** Comparison of a simulated three-state model (*k*_off,1_ = 0.01 frame^−1^, 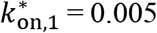 frame^−1^, *k*_off,2_ = 0.2 frame^−1^, 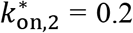 frame^−1^) with a predicted theoretical approximated two-species model, where the slower transitioning state is approximated as a separate immobile species (red) and the other species (blue) still contains two states with *k*_off,2_ and 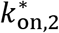 as transition rates. Upper panel track length of 4 steps, lower panel track length 8 steps. **(C)** Best fit of the simulated three-state model from (B) with a single-species two-state model. **(D)** MLE extraction of the expected fraction of the first approximated species for different values of *k*_off,2_ **(E)** Heat map of the log(*D_KS_*) between a simulated distribution (*D*_free_ = 1, fraction immobile = 0.1; ground truth (red dot)) and a theoretical predicted distributions with varying parameters around the parameters used for the simulation, where the simulation consisted either of 100.000 tracks at 10 ms frame time (left), 100.000 tracks at 50 ms frame time (middle) or 50.000 tracks at 10 ms frame time and 50.000 tracks at 50 ms frame time respectively (right). The Kolmogorov-Smirnov test statistic (*D_KS_*) is indicated in each histogram.

To test how well this approximation works and how well the method can distinguish this model from a simple two-state model, we simulated a linear (A ⇄ B ⇆ C) three-state model containing one slow transitioning bound state (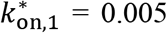 frame^−1^, *k*_off,1_ = 0.01 frame^−1^) and one fast transitioning bound state (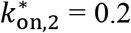 frame^−1^, *k*_off,2_ = 0.2 frame^−1^). We compared this simulation to our theoretically predicted distribution where we approximated the slower transitioning state as a separate immobile species and the faster transitioning state as a separate species (Figure 4B). The fraction of the approximated immobile species (20 %) and transitioning species (80 %) can be calculated from the ratio of the on- and off-rates (Figure 4B). We found very good agreement between the theoretical prediction and the simulation (*D_KS_* < 0.006) indicating that this approximation can be applied in this case. We then tried to find whether a single species two-state model could also fit the distribution of the three-state model (Figure 4C). We found that although for smaller track lengths there are parameters that can fit the distribution quite well (*D_KS_* = 0.0078 for track length of 4 steps), the distribution for larger tracks significantly deviated from the ground truth (*D_KS_* = 0.0149 for track length of 8 steps). Therefore, with a sufficient number of longer tracks two-state and three-state models are clearly distinguishable.

We then tested under which conditions the parameters can be reliably extracted from the data. To this end, we varied the transition rates of the fast-bound state (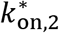 and *k*_off,2_) while keeping the slower bound state fixed. We observed that under all transition rates tested (0.1-10 transitions per frame), the error of the estimated parameters falls within 25% and that with increasing rates of the fast-bound state, the extraction of the fraction parameter became more reliable (Figure 4D). This finding indicates that as long as the transition rates associated with the different bound states are different enough (>10 fold), with one of them being significantly slower than the frame time used in the measurements, parameters for three state models can be reliably extracted with anaDDA.

More complex models with larger number of species, each having up to three states and meeting the requirements described above can also be fitted using anaDDA but are prone to increased uncertainty and under-/overfitting as many parameters in these models could give rise to similar distributions. To overcome this limitation, we implemented the ability to use data acquired at different frame times into a single global fit. By fitting data from multiple frame times simultaneously, the number of potential parameters that can fit all the data decreases, leading to more accurate and precise fitting.

As an example, we simulated a two-species (one immobile, one transitioning) model and calculated the Kolmogorov-Smirnov test statistic (*D_KS_*) for a range of parameters around the input parameters for a simulated dataset consisting of tracks either measured at a single frame time (10 or 50 ms) or a combined set where halve of the dataset contained simulated tracks from each frame time (Figure 4E). If there are other closely related parameters with similar *D_KS_* values to the ground truth, the fit can converge to these values as well. Therefore, the uncertainty is linked to the parameter space with *D_KS_* similar to the *D_KS_* of the ground truth. We observed that different frame times perform better on different parameters. In particular, short frame times led to more uncertainty in the determination of the fraction of each species, whereas long frame times gave more uncertainty in the determination of the free diffusion coefficient. When data recorded at different frame times is combined, there is only a single set of parameters that give rise to a similar *D_KS_* as the ground truth. In conclusion, the benefit of gathering data with different frame times is that it reduces the parameter space that can simultaneously fit multiple distributions and therefore offers better performance with the same number of data points.

### *E. coli* DNA polymerase I undergoes rapid DNA interactions

To test the applicability of our analysis method to experimental data, we re-analysed previously published data on the diffusion of DNA polymerase I in *E. coli*^17^. In this study, the diffusion distribution of PAmCherry-Pol1 was grouped into immobile and mobile diffusing particles by simple thresholding without determination of any transition kinetics. The authors found that under normal conditions only 4-5% of the proteins were immobile. However, they found that even the mobile tracks were mostly located within the nucleoid, which may suggest that these tracks represent transient DNA binding, probably probing the DNA for repair sites. We therefore hypothesized that the previously assigned mobile fraction is also undergoing rapid transitions between DNA bound and freely diffusing states.

We decided to fit the data with two species, one belonging to proteins involved in repair (a species with a single bound state) and one to probing (a species with a bound and a freely diffusing state). When we fitted this model (two species and three states; Figure 5A) we found a similar percentage of proteins involved in repair as described in the previous study (4%; Figure 5B). Furthermore, we found that the probing species had a free diffusion speed of 2.8 (± 0.2) μm^2^/s in the cytoplasm and that it is involved in vary rapid DNA probing events (*k*_off_ 137 ± 7 s^−1^; 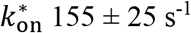). Based on the on and off-rates we calculated that the probing species spends more than half the time (~ 55%) bound to DNA. Altogether, DNA polymerase spends approximately ~ 60% bound to DNA either in repair (4%) or probing for mismatch sites (55%).

**Figure 5.**
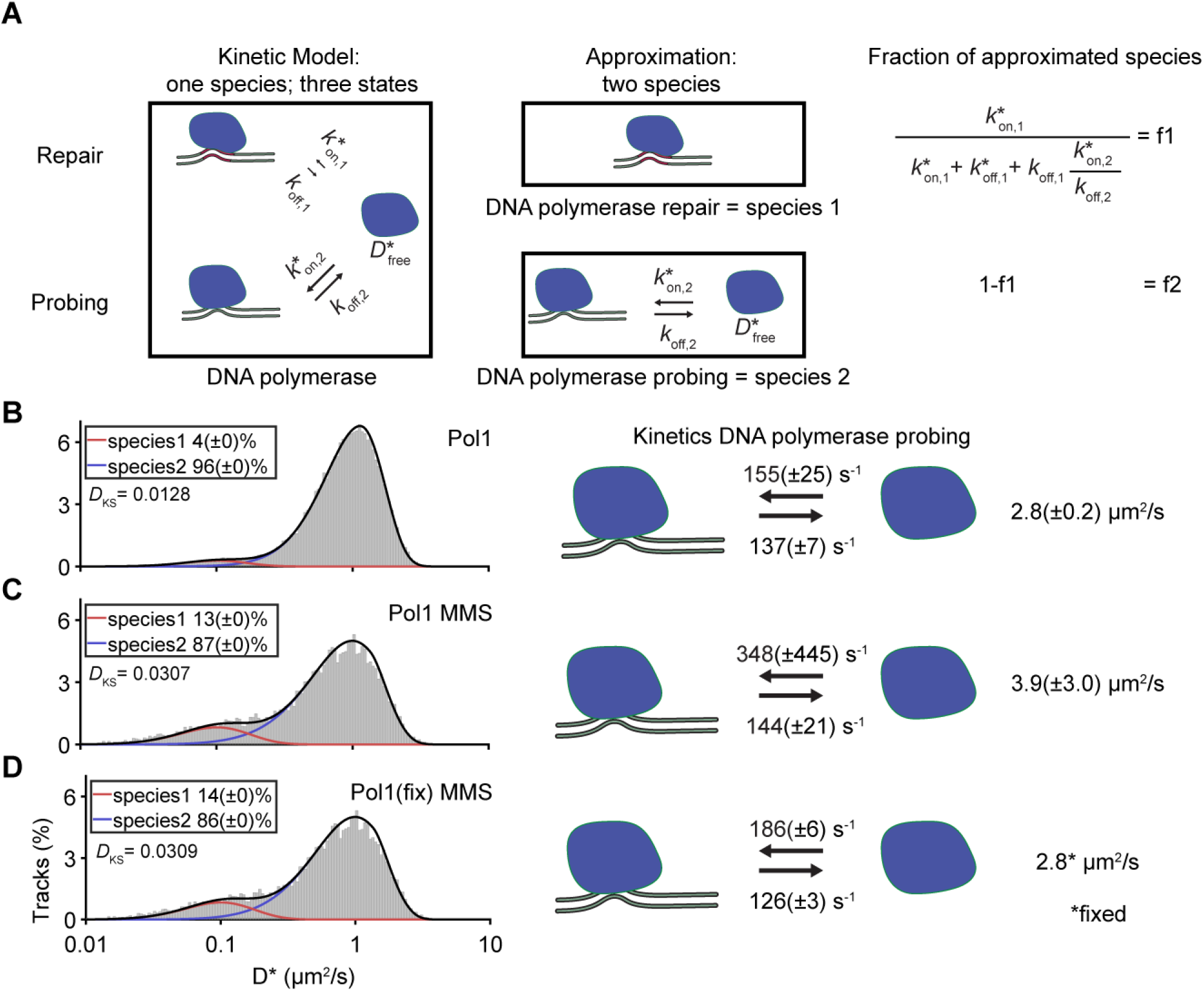
Extracting kinetic information of DNA polymerase I diffusing in live *E. coli*. **(A)** Approximated model of the kinetic model of DNA polymerase diffusion containing a DNA repair and DNA probing state (left). These states were separated into a single-state repair species (species 1; middle) and a probing species with two states (species 2; middle) The fraction of the two species are caused by the underlying ratios of the on- and off rates (right; Fig. 4A) **(B)** Fit of DNA polymerase I in untreated cells (n = 179.511 tracks). The *D*\* was fit with two species, one species involved in repair (red line) with a single state (immobile) and one species involved in scanning DNA with two states (mobile and immobile; blue line). The transition between the latter two states and the free diffusion coefficient of the mobile state are depicted. Fit was performed on all track lengths (1-8 steps) and histograms are only shown for a single-track length (4 steps). Tracks with more steps were truncated to 8 steps for the fit and 4 steps for the histogram. **(C)** Same as B but performed on data of DNA polymerase in cells treated with MMS. **(D)** Same as C except that the free diffusion coefficient of the mobile state was fixed to the same value as was found for polymerase in untreated cells (B).The Kolmogorov-Smirnov test statistic (*D_KS_*) is indicated in each histogram and uncertainty in parameters were estimated with bootstrapping (±SD). Experimental data was taken from a previous study^17^.

The study also measured the diffusivity of DNA polymerase in presence of the DNA damaging agent MMS. Using anaDDA, we found that the immobile species increased to 13% which matches the findings in the publication (13 ± 0.2%; Figure 5C). The transition rates and diffusion coefficients under this condition could not be assigned with confidence based on the bootstrap values (*k*_off_ 137 ± 7 s^−1^; 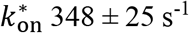). values. We hypothesized that this is caused by the lower number of available tracks (41.415 tracks) compared to the untreated dataset (142.178 tracks).

To quantitatively assess the transition kinetics in presence of DNA damage, we made the assumption that DNA damage would not alter the free diffusion behaviour of DNA polymerase in the cytoplasm but only the kinetics of the interactions with DNA. We therefore fixed *D*_free_ to the value found for DNA polymerase in untreated cells (2.8 μm^2^/s; Figure 5D) which caused the fitting to converge to a narrow range of transition rates. We observed that although the *k*_off_ remained the same (126 ± 3 s^−1^), the on-rate increased in the presence of damaged DNA (185 ± 6 s^−1^) indicating that more DNA polymerases were bound to DNA in long-term repair events (from 4 to 13%) and that also the polymerases engaged in probing spent more time bound to DNA. Altogether, these numbers would indicate that DNA polymerase in the presence of MMS spent ~ 75% of its time to DNA either at a repair site (13%) or while probing the DNA (60%).

We further found that the maximum step size of 5 pixels used in the original analysis significantly affected the distribution of observed *D*\* values (Figure S5). AnaDDA was able to correctly predict and take this effect into account. Overall, the transition rates between bound and unbound polymerase found under both conditions are high compared to the frame rate (>1 transition per frame), which demonstrates the applicability of anaDDA to quantify very fast transition kinetics *in vivo*.

## Discussion

Analytical diffusion distribution analysis (anaDDA) is able to accurately extract kinetics occurring within 4 orders of magnitude with around 10 to 0.01 transitions per frame. With conventional camera frame rates of 100 Hz, this range translates to interaction kinetics of 1 ms to 1 s even if the mean track length is as short as 3-4 frames. Furthermore, anaDDA is able to account for confinement and tracking window effects and has the possibility to fit data acquired at multiple frame times into a single global model. The re-analysis of previously published data on DNA polymerase I in *E. coli* suggests that this protein complex uses rapid probing of DNA and therefore spends more than 50% of its time bound to DNA, a value previously hypothesized based on its preferred localization in the nucleoid but not quantified up to now. These new insights into the biology of DNA polymerase *in vivo*, can experimentally be further tested. The predicted times spent on DNA in the absence (60%) and presence of MMS (75%) can be independently quantified by measuring the ratio of polymerases in DNA-containing and DNA-free segments of cells elongated by cephalexin as was done previously for CRISPR-Cas complexes in *E. coli*^47^.

Compared to other simulation-based frameworks for estimating transition rates^48–50^, anaDDA holds several advantages. First, the distributions of simulations are not exact as they are generated from a limited number of particles and therefore do not allow for using an MLE approach, which requires convergence based on exact probability even for small changes in the parameter space. Secondly, since analysis methods can only be verified by knowing the ground truth, these algorithms can only be tested with and against simulations itself. Consequently, the analysis and verification data are not independent, which could lead to unobservable errors. Furthermore, our analysis method is computationally significantly faster. MLE takes just around 15 s to find the optimal parameter set for a global fit to a 50.000 tracks dataset with a track length range of 1-8 steps (Intel Core i7), whereas a simulation estimating three parameters with a global fit of all step sizes, required around 10 hours to find an optimal set of parameters.

So far, it is possible to include two transitioning state into the direct fitting. We have shown, however, that when transition rates are slow compared to the frame time of the measurement, states can be treated as separate species. Further development of the underlying master equations of PDA statistics could allow direct implementation of multistate models.

With the increasing use of brighter and more stable organic fluorophore^14,51^ or low photon flux measurements^52^ for single-particle tracking, the resulting increase of the track length and the decrease of the localization error will enable further improvements in the precision of extracted kinetic parameters. Currently, we have implemented the software for tracking in two dimensions, but the algorithms can be further modified towards tracking in three dimensions. Using the estimated error for each individual localization can further improve the robustness of the analysis as has been demonstrated previously^53^. Another improvement which can be incorporated in our framework and has already been developed is to take the effect of particles moving out-of-focus, and the recovery of localizations depending on diffusion coefficients into account^38,50^.

Our analysis method allows the quantification of fast kinetic transitions inside living cells with state lifetimes in the 1 ms to 1 s range opening a temporal range at which many DNA screening interactions are expected to take place^54^. So far however, quantifying these interactions has been limited due to a lack of appropriate analytic and experimental methods. We are convinced that anaDDA will offer the means to determining fast kinetics *in vivo* which will be key to uncover and understand the behaviour of biomolecular complexes in cells.

## Acknowledgements

The authors thank Dr. S. Uphoff for supplying the experimental data on DNA polymerase and K. Martens for feedback and co-development of the Monte Carlo DDA implementation (MC-DDA) and all members of the Hohlbein and the Brouns groups for input during group discussions. This work was supported by the European Research Council (ERC) Stg grant 639707 and by the Netherlands Organisation for Scientific Research (NWO/OCW), as part of the Frontiers of Nanoscience (NanoFront) program.

## Software

The latest version of the software is available on GitHub: https://github.com/HohlbeinLab/anaDDA

## Author contributions

S.B. and J.H. conceived and supervised the project; J.H. provided the initial idea, J.V. developed the framework, derived the equations and wrote the analysis scripts, J.V. and J.H. wrote the manuscript and S.B. provided feedback on the manuscript.

## References

1. Miller, H., Zhou, Z., Shepherd, J., Wollman, A. J. M. & Leake, M. C. Single-molecule techniques in biophysics: A review of the progress in methods and applications. Reports on Progress in Physics 81, (2018).

2. Hohlbein, J. et al. Conformational landscapes of DNA polymerase I and mutator derivatives establish fidelity checkpoints for nucleotide insertion. Nat. Commun. 4, 2131 (2013).

3. Hodges, C., Bintu, L., Lubkowska, L., Kashlev, M. & Bustamante, C. Nucleosomal fluctuations govern the transcription dynamics of RNA polymerase II. Science 325, 626–628 (2009).

4. Rothenberg, E. & Ha, T. Single-molecule FRET analysis of helicase functions. Methods Mol. Biol. 587, 29–43 (2010).

5. Craig, J. M. et al. Revealing dynamics of helicase translocation on single-stranded DNA using high-resolution nanopore tweezers. Proc. Natl. Acad. Sci. U. S. A. 114, 11932–11937 (2017).

6. Rutkauskas, M., Krivoy, A., Szczelkun, M. D., Rouillon, C. & Seidel, R. in Methods in Enzymology 582, 239–273 (2017).

7. Blosser, T. R. et al. Two distinct DNA binding modes guide dual roles of a CRISPR-Cas protein complex. Mol. Cell 58, 60–70 (2015).

8. Heller, I., Hoekstra, T. P., King, G. A., Peterman, E. J. G. & Wuite, G. J. L. Optical tweezers analysis of DNA-protein complexes. Chemical Reviews 114, 3087–3119 (2014).

9. Blouin, S., Craggs, T. D., Lafontaine, D. A. & Penedo, J. C. in Methods in Molecular Biology 1334, 115–141 (2015).

10. Hohlbein, J., Craggs, T. D. & Cordes, T. Alternating-laser excitation: Single-molecule FRET and beyond. Chemical Society Reviews 43, 1156–1171 (2014).

11. Lerner, E. et al. Toward dynamic structural biology: Two decades of single-molecule förster resonance energy transfer. Science (New York, N.Y.) 359, (2018).

12. Kapanidis, A. N., Lepore, A. & El Karoui, M. Rediscovering Bacteria through Single-Molecule Imaging in Living Cells. Biophysical Journal 115, 190–202 (2018).

13. Jradi, F. M. & Lavis, L. D. Chemistry of Photosensitive Fluorophores for Single-Molecule Localization Microscopy. ACS Chemical Biology 14, 1077–1090 (2019).

14. Banaz, N., Mäkelä, J. & Uphoff, S. Choosing the right label for single-molecule tracking in live bacteria: Side-by-side comparison of photoactivatable fluorescent protein and Halo tag dyes. J. Phys. D. Appl. Phys. (2018). doi:10.1088/1361-6463/aaf255

15. Manley, S. et al. High-density mapping of single-molecule trajectories with photoactivated localization microscopy. Nat. Methods 5, 155–157 (2008).

16. English, B. P. et al. Single-molecule investigations of the stringent response machinery in living bacterial cells. Proc. Natl. Acad. Sci. 108, 365–373 (2011).

17. Uphoff, S., Reyes-Lamothe, R., Garza de Leon, F., Sherratt, D. J. & Kapanidis, A. N. Single-molecule DNA repair in live bacteria. Proc. Natl. Acad. Sci. U. S. A. 110, 8063–8068 (2013).

18. Garza de Leon, F., Sellars, L., Stracy, M., Busby, S. J. W. & Kapanidis, A. N. Tracking Low-Copy Transcription Factors in Living Bacteria: The Case of the lac Repressor. Biophys. J. 112, 1316–1327 (2017).

19. Van Beljouw, S. P. B. et al. Evaluating single-particle tracking by photo-activation localization microscopy (sptPALM) in Lactococcus lactis. Phys. Biol. (2019). doi:10.1088/1478-3975/ab0162

20. Qian, H., Sheetz, M. P. & Elson, E. L. Single particle tracking. Analysis of diffusion and flow in two-dimensional systems. Biophys. J. 60, 910–921 (1991).

21. Saxton, M. J. Single-particle tracking: The distribution of diffusion coefficients. Biophys. J. 72, 1744–1753 (1997).

22. Kalinin, S., Felekyan, S., Valeri, A. & Seidel, C. A. M. Characterizing Multiple Molecular States in Single-Molecule Multiparameter Fluorescence Detection by Probability Distribution Analysis. J. Phys. Chem. B 112, 8361–8374 (2008).

23. Palo, K., Mets, Ü., Loorits, V. & Kask, P. Calculation of photon-count number distributions via master equations. Biophys. J. 90, 2179–2191 (2006).

24. Nir, E. et al. Shot-noise limited single-molecule FRET histograms: Comparison between theory and experiments. J. Phys. Chem. B 110, 22103–22124 (2006).

25. Santoso, Y., Torella, J. P. & Kapanidis, A. N. Characterizing Single-Molecule FRET Dynamics with Probability Distribution Analysis. ChemPhysChem 11, 2209–2219 (2010).

26. Farooq, S. & Hohlbein, J. Camera-based single-molecule FRET detection with improved time resolution. Phys. Chem. Chem. Phys. 17, 27862–27872 (2015).

27. Persson, F., Lindén, M., Unoson, C. & Elf, J. Extracting intracellular diffusive states and transition rates from single-molecule tracking data. Nat. Methods 10, 265–9 (2013).

28. Karslake, J. D. et al. SMAUG: Analyzing single-molecule tracks with nonparametric Bayesian statistics. bioRxiv (2019). doi:10.1101/578567

29. Plank, M., Wadhams, G. H. & Leake, M. C. Millisecond timescale slimfield imaging and automated quantification of single fluorescent protein molecules for use in probing complex biological processes. Integr. Biol. (2009). doi:10.1039/b907837a

30. Stracy, M. et al. Live-cell superresolution microscopy reveals the organization of RNA polymerase in the bacterial nucleoid. Proc. Natl. Acad. Sci. 112, E4390–E4399 (2015).

31. Vrljic, M., Nishimura, S. Y., Brasselet, S., Moerner, W. E. & McConnell, H. M. Translational diffusion of individual class II MHC membrane proteins in cells. Biophys. J. 83, 2681–2692 (2002).

32. Michalet, X. Mean square displacement analysis of single-particle trajectories with localization error: Brownian motion in an isotropic medium. Phys. Rev. E - Stat. Nonlinear, Soft Matter Phys. 82, 041914 (2010).

33. Antonik, M., Felekyan, S., Gaiduk, A. & Seidel, C. A. M. Separating structural heterogeneities from stochastic variations in fluorescence resonance energy transfer distributions via photon distribution analysis. J. Phys. Chem. B 110, 6970–6978 (2006).

34. Berglund, A. J. Statistics of camera-based single-particle tracking. Phys. Rev. E - Stat. Nonlinear, Soft Matter Phys. 82, (2010).

35. Paris, J. F. A Note on the Sum of Correlated Gamma Random Variables. Available in arxiv.org at http://arxiv.org/abs/1103.0505. (2011). at <http://arxiv.org/abs/1103.0505>

36. Martos-Naya, E., Romero-Jerez, J. M., Lopez-Martinez, F. J. & Paris, J. F. A MATLAB TM program for the computation of the confluent hypergeometric function Φ 2.

37. Lee, B. H. & Park, H. Y. HybTrack: A hybrid single particle tracking software using manual and automatic detection of dim signals. Sci. Rep. 8, (2018).

38. Hansen, A. S. et al. Robust model-based analysis of single-particle tracking experiments with Spot-On. Elife 7, (2018).

39. Uphoff, S., Sherratt, D. J. & Kapanidis, A. N. Visualizing protein-DNA interactions in live bacterial cells using photoactivated single-molecule tracking. J. Vis. Exp. (2014). doi:10.3791/51177

40. Bickel, T. A note on confined diffusion. Phys. A Stat. Mech. its Appl. 377, 24–32 (2007).

41. Bausch, J. On the efficient calculation of a linear combination of chi-square random variables with an application in counting string vacua. J. Phys. A Math. Theor. (2013). doi:10.1088/1751-8113/46/50/505202

42. Smith, S. W. in Digital Signal Processing 311–318 (2003). doi:10.1016/b978-0-7506-7444-7/50055-8

43. Myung, I. J. Tutorial on maximum likelihood estimation. J. Math. Psychol. 47, 90–100 (2003).

44. Vink, J. N. A. et al. Direct Visualization of Native CRISPR Target Search in Live Bacteria Reveals Cascade DNA Surveillance Mechanism. Mol. Cell 589119 (2019). doi:10.1016/j.molcel.2019.10.021

45. Woodside, M. T. et al. Direct measurement of the full, sequence-dependent folding landscape of a nucleic acid. Science 314, 1001–1004 (2006).

46. Slutsky, M. & Mirny, L. A. Kinetics of Protein-DNA Interaction: Facilitated Target Location in Sequence-Dependent Potential. Biophys. J. 87, 4021–4035 (2004).

47. Vink, J. N. A. et al. Direct Visualization of Native CRISPR Target Search in Live Bacteria Reveals Cascade DNA Surveillance Mechanism. Mol. Cell (2020). doi:10.1016/j.molcel.2019.10.021

48. Wieser, S., Axmann, M. & Schütz, G. J. Versatile analysis of single-molecule tracking data by comprehensive testing against Monte Carlo simulations. Biophys. J. 95, 5988–6001 (2008).

49. Martens, K. J. A. et al. Visualisation of dCas9 target search in vivo using an open-microscopy framework. Nat. Commun. 10, 3552 (2019).

50. Rocha, J., Corbitt, J., Yan, T., Richardson, C. & Gahlmann, A. Resolving Cytosolic Diffusive States in Bacteria by Single-Molecule Tracking. Biophys. J. 116, 1970–1983 (2019).

51. Los, G. V. et al. HaloTag: A novel protein labeling technology for cell imaging and protein analysis. ACS Chem. Biol. 3, 373–382 (2008).

52. Balzarotti, F. et al. Nanometer resolution imaging and tracking of fluorescent molecules with minimal photon fluxes. Science 355, 606–612 (2017).

53. Lindén, M. & Elf, J. Variational Algorithms for Analyzing Noisy Multistate Diffusion Trajectories. Biophys. J. (2018). doi:10.1016/j.bpj.2018.05.027

54. Elf, J., Li, G. W. & Xie, X. S. Probing transcription factor dynamics at the single-molecule level in a living cell. Science 316, 1191–1194 (2007).

