## Supplemental Figures for "Extracting transition rates in single-particle tracking using analytical diffusion distribution analysis"

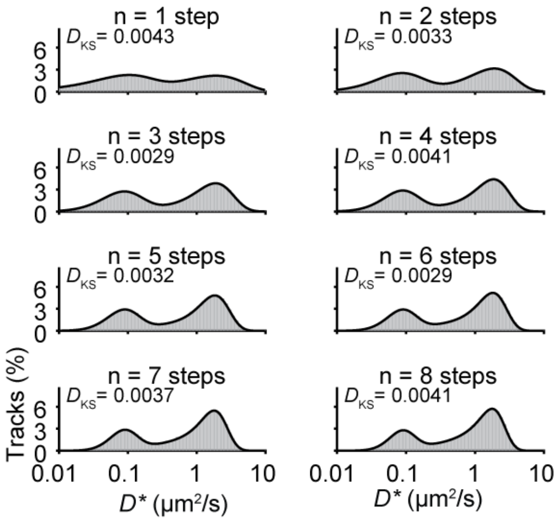


**Figure S1.** Diffusion distributions for different number of steps within a single trajectory, simulated (grey boxes) and DDA predicted distributions (black line) for 1-8 number of steps. Simulation parameters: $k^*_{\mathrm{on}}$ = 0.2 frame$^{-1}$, $k^*_{\mathrm{on}}$ = 0.2 frame$^{-1}$, $D_{\mathrm{free}}$ = 4 μm$^2$/s, σ = 30 nm (localisation precision) and N = 50.000. The Kolmogorov-Smirnov test statistic ($D_{KS}$) is indicated at each histogram.

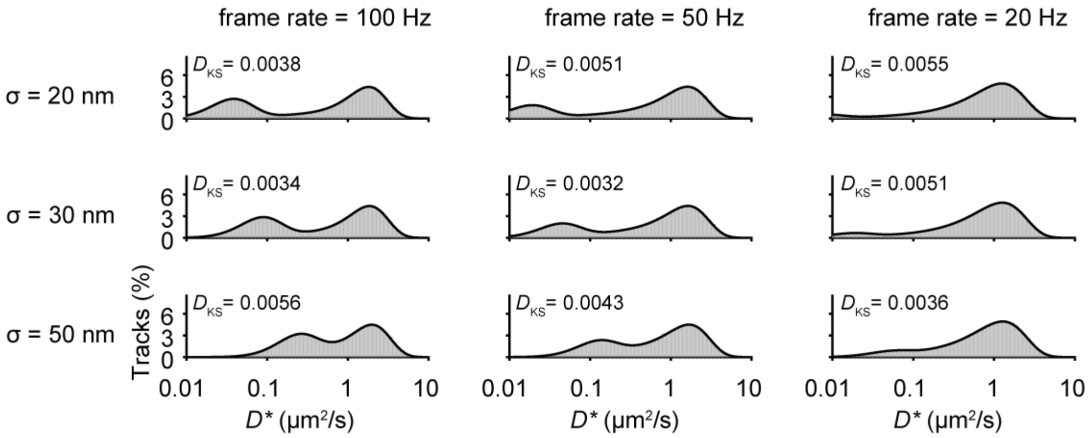


**Figure S2.** The effect of framerate and localization error on the shape of the distributions. The predicted probability distribution of analytical DDA (black) closely resembles the simulated values (grey). Increasing the framerate shifts the peak of the bound population left to lower *D** values and increased localization errors shift this peak right to higher *D** values. Simulation parameters: $k^*_{\mathrm{on}}$ = 0.02 frame$^{-1}$, $k^*_{\mathrm{on}}$ = 0.02 frame$^{-1}$, $D_{\mathrm{free}}$ = 4 μm$^2$/s and N = 50.000. The Kolmogorov-Smirnov test statistic ($D_{KS}$) is indicated at each histogram.

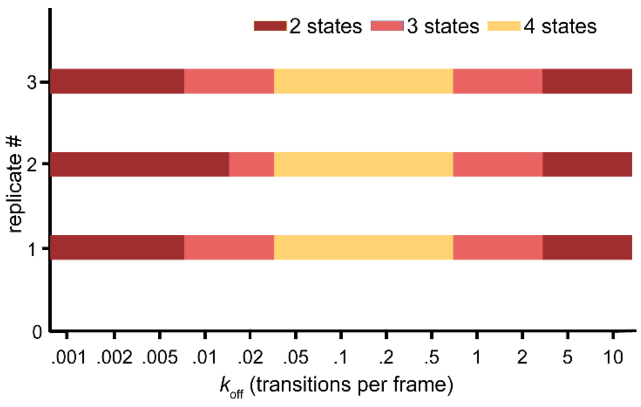


**Figure S3.** The number of states extracted from vbSPT in a simulated two-state system. After removing restrictions on the maximum amount of states in vbSPT, the number of states fitted under some conditions differed from the amount of states modelled (2 states). Indicated are which simulated replicates contained which number of states (dark red: 2 states, light red: 3 states and yellow 4 states. Other parameters included were: $k^{*}_{\mathrm{on}}$ = $k_{\mathrm{off}}$, $D_{\mathrm{free}}$ = 4 μm$^2$/s and σ = 30 nm.

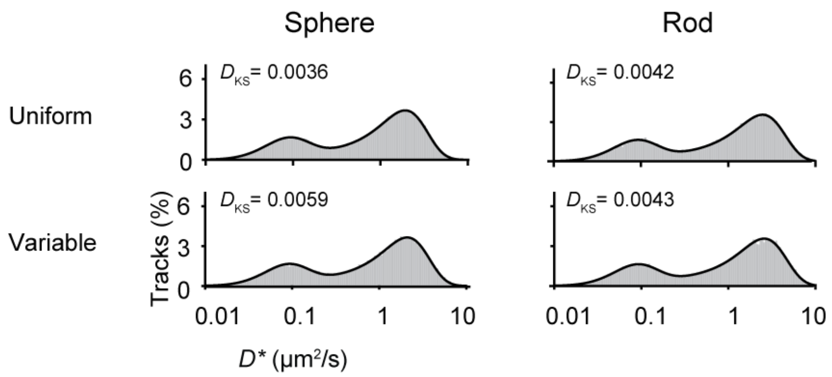


**Figure S4. Distributions of a population of cells with uniform and variable cell size.** The cell shapes were either spherical (left) or rod-shaped (right; radius to length ratio is 1:8). The average radius for uniform (upper row) and variable (lower row) cell sizes was the same: $r_{\text{confined}} = \sqrt{5D_{\text{free}}t}$. For the variable cell size, 60% of the cells were simulated with the same size as the average, whereas for 20% of the cells were simulated 25% smaller and for 20% of the cells were simulated with 25% larger cells. Further parameters used were: $k^*_{\text{on}}$= 0.02 frame$^{-1}$, $k^*_{\text{on}}$ = 0.02 frame$^{-1}$, $D_{\text{free}}$ = 4 μm$^2$/s, σ = 30 nm and $dt$ = 0.01 s. The Kolmogorov-Smirnov test statistic ($D_{KS}$) is indicated at each histogram.

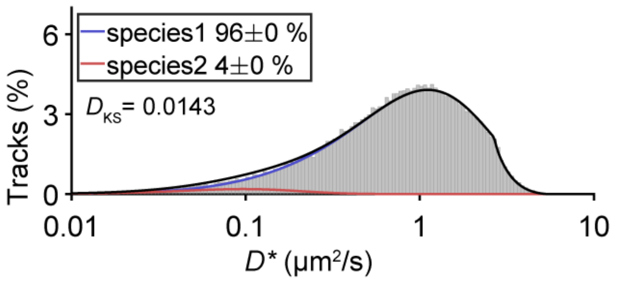


**Figure S5. DNA polymerase histogram for tracks with a track length of 2 steps.** The same condition and fitting parameters as for Figure 5A were used except that a two-step track length is shown here. The maximum step size 5 pixels (0.6 μm) that was initially applied to this dataset results in a discontinuous distribution, which is correctly captured by the ana-DDA fit. Data from previous study[17].
